# *Mycobacterium smegmatis* expands across surfaces using hydraulic sliding

**DOI:** 10.1101/2020.09.22.307801

**Authors:** Eric J.G. Pollitt, Oliver Carnell, Egbert Hoiczyk, Jeffrey Green

**Author notes:** Corresponding author Correspondence: Eric J.G. Pollitt, Biomedical Science, University of Sheffield, Firth Court, Western Bank, Sheffield, S10 2TN.

## Abstract

*Mycobacterium smegmatis* spreads over soft agar surfaces by sliding motility, a form of passive motility in which growth and reduction of surface adhesion enable the bacteria to push each other outwards. Hence, sliding motility is mostly associated with round colonies. However, *M. smegmatis* sliding colonies can also produce long, pointed dendrites. Round sliding colonies were readily reproduced, but our non-round colonies were different from those seen previously. The latter (named digitate colonies) had centimetre-long linear protrusions, containing a central channel filled with a free-flowing suspension of *M. smegmatis* and solid aggregates. Digitate colonies had both a surface pellicle and an inner biofilm component surrounding a central channel, which sat in a cleft in the agar. Time-lapse microscopy showed that the expansion of the fluid-filled channel enabled the lengthwise extension of the protrusions without any perceptible growth of the bacteria taking place. These observations represent a novel type of sliding motility, named hydraulic sliding, associated with a specialised colony structure and the apparent generation of force by expansion of a liquid core. As this structure requires pellicle formation without an initial liquid culture it implies the presence of an unstudied mycobacterial behaviour that may be important for colonisation and virulence.

**Originality-Significance Statement:** This study is the first to identify a new form of passive motility in the mycobacteria; hydraulic sliding, in which liquid expansion is the cause of motility. This form of motility has so far never been described in bacteria. The study also reveals new ways mycobacteria can form biofilms and colonize complex three-dimensional substrates, aspects of mycobacterial biology that are important for infection, pathogenesis and vaccine development.

**Author Summary:** *Mycobacterium smegmatis* is used as a non-pathogenic model organism for pathogenic mycobacteria. During growth, *M. smegmatis* can move passively over soft agar surfaces by a process called sliding motility, whereby colony growth directly pushes cells outwards. Although passive, sliding motility is believed to be important in allowing bacteria to colonise surfaces. Sliding motility however does not fully account for how *M. smegmatis* produces dendritic colonies. We attempted to generate dendritic colonies but found instead that the cells produced colonies that had larger protrusions radiating from them (digitate colonies). Digitate colonies are a previously unobserved phenomenon, in that the bacteria create a biofilm-lined, fluid-filled, pellicle-covered, deep cleft in the agar and move across the surface by the expansion of the contained liquid core of the protrusions. Given the new structure and the new mechanism of expansion we have termed this set of behaviours hydraulic sliding. These observations are important as it is a new variation in the way bacteria can move, generate biofilms (notably mycobacterial pellicle) and colonize complex three-dimensional substrates.

## Introduction

*M. smegmatis* is a non-motile bacterium which, due to its relatively fast growth, is used as a model for other mycobacteria, particularly *Mycobacterium tuberculosis*. Although it is non-motile, it is able to move passively across soft media by a process called sliding (Martínez *et al*., 1999). Sliding motility is a form of passive motility that was originally defined by Henrichsen as “a kind of surface translocation produced by expansive forces in a growing culture in combination with special surface properties” (Henrichsen, 1972). Thus, sliding is dependent on bacterial growth pushing cells outwards facilitated by factors that reduce surface adhesion (Hölscher and Kovács, 2017). Sliding motility has been reported for several bacteria; some move by this mechanism alone (such as *M. smegmatis* and *Streptococcus spp*.) whereas others move in this way if other motility mechanisms have been disabled (e.g. *Pseudomonas aeruginosa* and *Legionella pneumophillia*), or under special growth/surface conditions (e.g. *Bacillus subtilis* and *Staphylococcus aureus*), or a combination of the above (Henrichsen, 1972; Murray and Kazmierczak, 2008; Shrout, 2015; Hölscher and Kovács, 2017; Kovács *et al*., 2017). Thus, sliding is believed to be a means by which bacteria move across surfaces in the absence of active motility mechanisms. Interestingly, the factors that reduce surface adhesion to allow sliding to occur have frequently proved to be important virulence factors, such as the Phenol Soluble Modulins (PSMs) in *S. aureus*, and rhamnolipids in *P. aeruginosa* (Kaito and Sekimizu, 2007; Hölscher and Kovács, 2017).

*M. smegmatis* sliding was first reported by Martínez *et al*. (Martínez *et al*., 1999). They found that *M. smegmatis* strain mc^2^155 grown on 7H9 medium solidified with Difco agar produced colonies with long, thin dendrites. However, when agarose was used as a solidifying agent, circular round colonies were observed. For both sliding colony morphotypes the edges consisted of a monolayer of cells that moved outwards as a compact group. This movement was linked to growth of the colony, lacked behaviours associated with active motility, and hence was assigned as sliding motility (Martínez *et al*., 1999). The cells were also organised as what were termed ‘pseudofilaments’, where the separate bacteria are linked at or near their poles in small branching chains of <20 bacteria. This is fundamentally different from the individual cells/disorganised clumps observed in liquid culture. All subsequent studies of the genetic basis of *M. smegmatis* sliding have used the agarose based motility assay that generates round colonies (Recht *et al*., 2000; Recht and Kolter, 2001; Gopalaswamy *et al*., 2008; Jamet *et al*., 2015). It has been found that sliding in these circumstances is primarily dependent on glycopeptidolipids (GPLs), with mutations that affect sliding being associated with GPL production (Martínez *et al*., 1999; Recht *et al*., 2000). Glycopeptidolipids are found in the cell envelope of the non-tuberculous mycobacteria (Schorey and Sweet, 2008; Mukherjee and Chatterji, 2012). Disruption of GPL production alters colony morphology, which is known to be important in biofilm formation and is likely to be implicated in virulence and drug susceptibility in pathogenic mycobacteria (Schorey and Sweet, 2008; Chakraborty and Kumar, 2019). Furthermore, even though GPLs are not covalently bound to the mycobacterial cell envelope, they are unlikely to freely diffuse around the colony, because movement of a non GPL-producing strain was not restored by GPL producers, no surfactants were detected by the droplet collapse method, and exogenous surfactant did not restore motility to non-GPL strains (Jain *et al*., 1991; Recht *et al*., 2000). It is quite common for bacteria to secrete free surfactant to move across surfaces, but some forms of sliding have been proposed to occur without surfactants, and to date no surfactants have been shown to be involved in *M. smegmatis* sliding (Kearns, 2010; Hölscher and Kovács, 2017).

In comparison to *M. smegmatis*, where the long thin dendrites were associated with sliding motility, *S. aureus* colonies can also engage in sliding motility and also produce dendrites, but rather than being generated through sliding motility the dendrites resulted from motile gliding comets (Martínez *et al*., 1999; Pollitt *et al*., 2015; Pollitt and Diggle, 2017). *S. aureus* sliding motility requires cell surface teichoic acids and a diffusible surfactant component, the PSMs, which are secreted by an essential translocase (Kaito and Sekimizu, 2007; Tsompanidou *et al*., 2013; Otto, 2014). The differences in dendrite production by *M. smegmatis* and *S. aureus*, in which the former are generated by sliding motility and the latter are not, prompted further investigation of the unusual characteristics associated with the sliding motility of *M. smegmatis*. Sliding is not usually associated with long, thin dendrites (Henrichsen, 1972; Hölscher and Kovács, 2017; Pollitt and Diggle, 2017). As noted above, Henrichsen previously identified that sliding motility should be associated with round colonies as the bacteria are passively pushing each other out in all directions and inherently have no control over direction (Henrichsen, 1972). Indeed, most bacteria that exclusively engage in sliding motility produce round or frond-like colonies (the fronds likely forming due to differences in growth rate). Hence, *M. smegmatis* is unusual in forming long, thin, pointed dendrites, whilst engaging in sliding motility (Hölscher and Kovács, 2017). Here, we report a new colony morphology for *M. smegmatis* (digitate colonies) displaying centimetre-long protrusions containing a free-flowing suspension of bacteria covered by a pellicle. Experiments suggested that fluid accumulation enabled the expansion of the protrusions and spread of the colony without visible growth, representing a new form of sliding motility, which we term hydraulic sliding.

## Results

### *Observation of a new* Mycobacterium smegmatis *colony morphology (digitate colonies)*

The starting point for this work was the report from Martínez *et al*. demonstrating that *M. smegmatis* sliding motility was associated with at least two colony forms, i.e. circular colonies and colonies with long, thin dendrites (Martínez *et al*., 1999). Here, the circular sliding colonies were readily observed on agarose medium (Fig. 1a). However, it was not possible to consistently replicate the dendrite morphotype using Bacto agar as a solidifying agent. Very occasionally short branching dendrites were observed, but the long dendrites previously reported were not formed. When we replaced Bacto agar with Noble agar, colonies with centimetre-long protrusions radiating from the centre that were much broader and straighter than those previously reported, were observed (higher agar concentrations prevented their formation, Fig. 1b-d). Remarkably the protrusions readily climbed the meniscus of the agar and could extend upwards at a steep angle (Fig. 1e). To distinguish this new morphotype from those previously described (Martínez *et al*., 1999), we henceforth refer to them as digitate colonies.

**Fig. 1.**
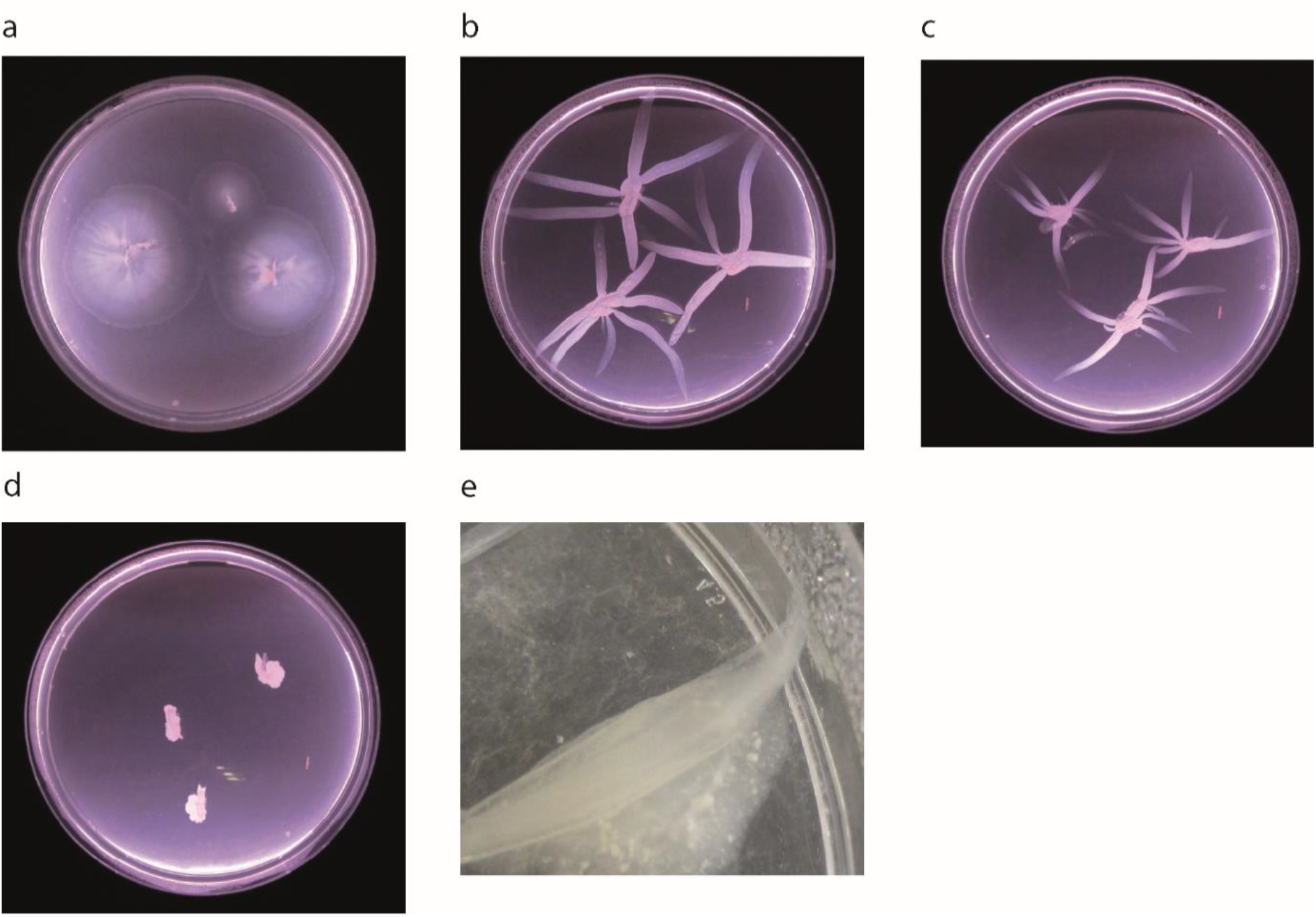
*Mycobacterium smegmatis* colony morphotypes. (a) Circular colonies on agarose medium 3 days post-inoculation; (b-d) digitate colonies on: (b) 0.2% Noble agar medium; (c) 0.25% Noble agar; and (d) 0.3% Noble agar 3 days post-inoculation. (e) Protrusion of a digitate colony extending directly up the meniscus.

### *Digitate colonies of* Mycobacterium smegmatis *have liquid filled protrusions*

The digitate colonies formed on Noble agar were different from any previously reported *M. smegmatis* colonies (Martínez *et al*., 1999). Developing over 3 days, on average, 3-5 broad protrusions emanated from each colony (Fig. 2a-c). The protrusions differed from the previously observed dendritic colony morphology in that: 1) each protrusion was linear with no branching; 2) the protrusions were broader; and 3) within each protrusion there was a core of mobile liquid containing suspended bacteria and large aggregates. The liquid core extended almost to the tip of the protrusion and was fully mobile within the protrusion, but was retained when the plate was briefly inverted or rocked (Fig. 3; Video S1). Introduction of Phenol Red to the core at the tip of a protrusion followed by rocking the plate showed that the dye flowed rapidly to the other tips, indicating that the protrusion cores are connected and the liquid moved from one protrusion to another (Fig. 3). The protrusions did not avoid other protrusions and occasionally merged.

**Fig. 2.**
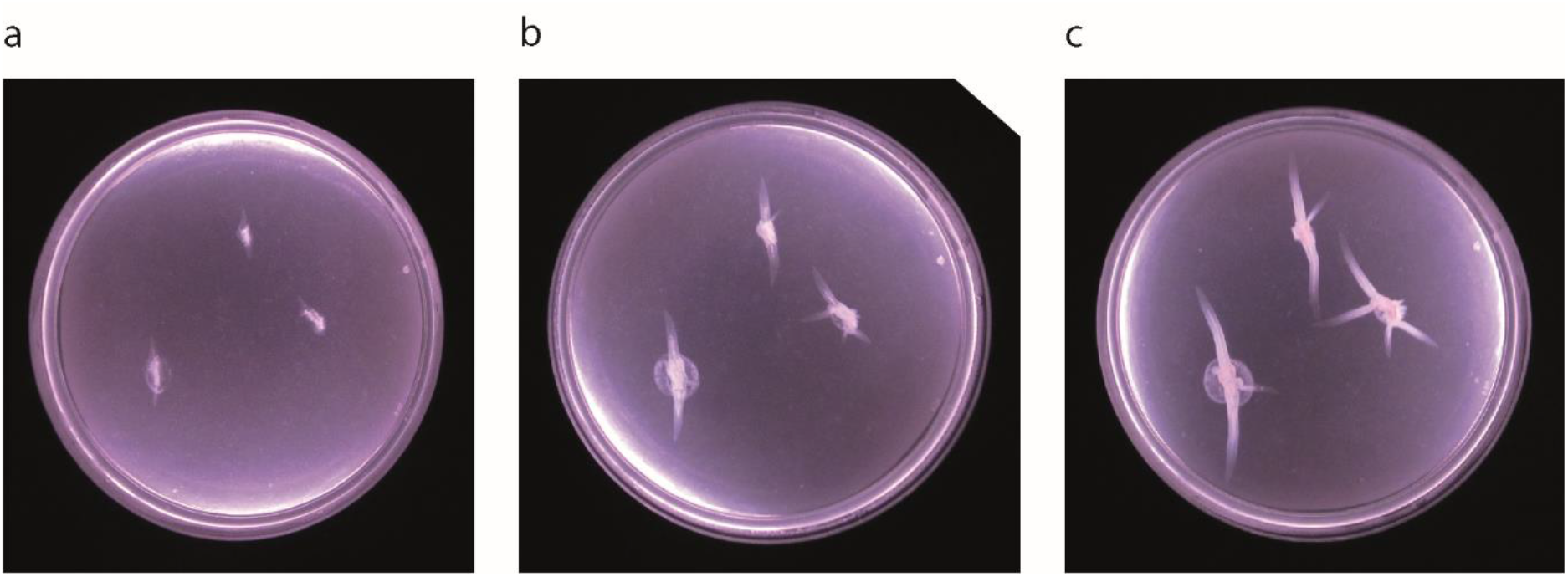
Formation of *M. smegmatis* digitate colonies with broad, linear protrusions. The agar plate was inoculated at three points with *M. smegmatis* and incubated at 37°C. The plate was imaged: (a) 24 h; (b) 48 h; and (c) 72 h, post-inoculation.

**Fig. 3.**
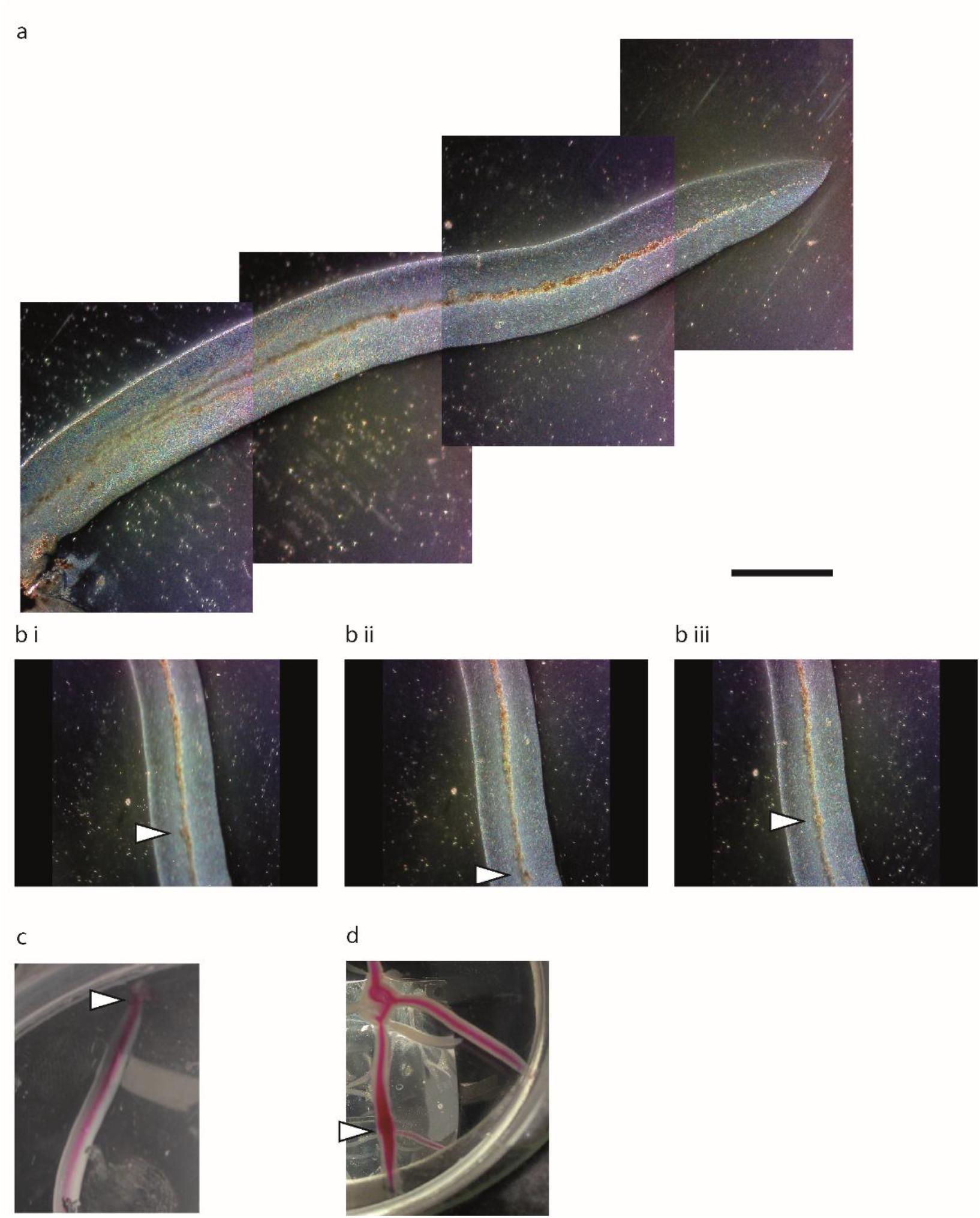
Dissecting microscope imaging of an *M. smegmatis* digitate colony. (a) A protrusion imaged using a dissecting microscope (x4 magnification), note the central channel within. Scale Bar: 5 mm (b) Selected sequential images (i-iii) from Video S1. As the plate is rocked the aggregates move within the central core, the large aggregate indicated by the arrows is tracked up and down the core (see arrows and Video S1). Nothing outside the central channel moves. (c) Injection of phenol red into the central core. (i) Spotting of 1 μl of phenol red (10%) at the tip and then tilting the plate causes the dye to follow the central core; (ii) spotting 5 μl of phenol red and tilting the plate allows the dye to move through the core to its maximum extent, spreading from one protrusion to another within the digitate colony. (Arrow indicates site of injection).

Examination of the tips and edges of the protrusions by light microscopy revealed the presence of mycobacterial pseudofilaments arranged as branching chains of less than 20 individuals (Fig. 4a). These did not occur in liquid cultures, where single and small aggregates of bacteria were seen (Fig. 4b), but were a feature of the circular colonies (Fig. 4c). These branching pseudofilaments have been previously identified by Martínez *et al*. (1999). In the digitate colonies, the bacteria broadly aligned with the edge of the protrusion and curved around at the tip, whereas the bacteria in the circular colonies were not aligned in any direction.

**Fig. 4.**
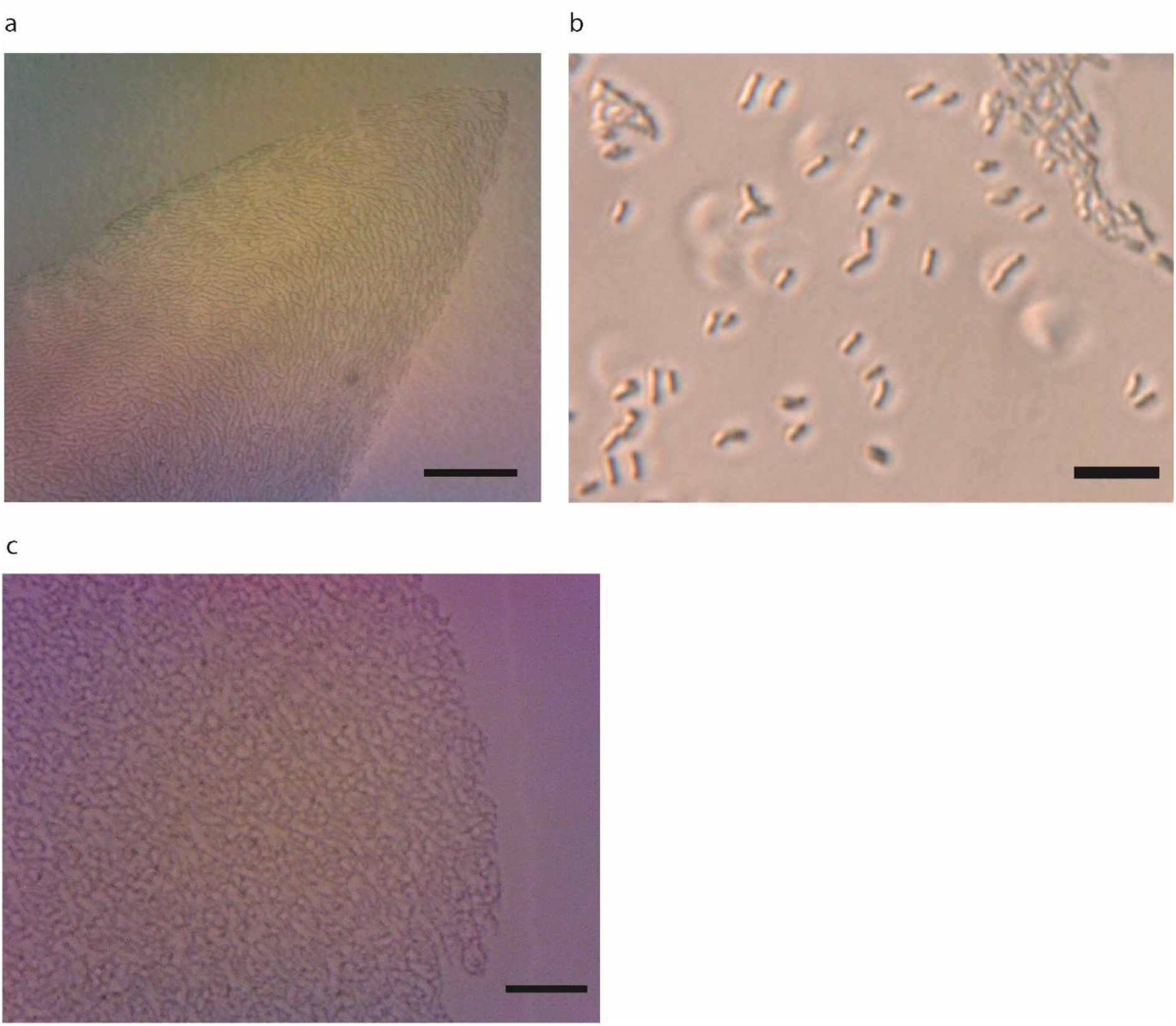
Light microscopy of sliding colonies. (a) The tip of an *M. smegmatis* digitate colony. The bacteria are arranged as branching pseudofilaments which broadly align with the curve of the tip. Scale Bar: 100 µm. (b) Image of *M. smegmatis* cells grown in 7H9 liquid culture on a glass slide showing that the cells do not branch. Scale Bar: 20 µm (c) The edge of a circular *M. smegmatis* colony also showing unaligned branching pseudofilaments. Scale Bar: 50 µm.

Specimens were flattened and observed under a light microscope to investigate the nature of the upper surface of the protrusions. The top of the protrusion was preserved and appeared to be a comparatively thick (at least 20 cell layers) pellicle that resembled the mycobacterial pellicles generated on the surface of liquid cultures (Fig. 5; Pang *et al*., 2012; Parvez *et al*., 2018). The bacteria were inoculated at a solid-air interface and proceeded to create a liquid-air interface biofilm; a phenomenon apparently not previously observed. The top layer of the pellicle had a different consistency from the edges of the colony indicating that they differ; the edges are likely to be more strongly attached to the agar and could tear away from the main pellicle when the structure was disrupted. We also noted that wrinkling, as happens with mature *M. smegmatis* pellicle grown on liquid culture, was not observed (Ojha *et al*., 2005). In contrast when the circular colonies were flattened, they were reduced to a liquid suspension with no perceptible structures. To investigate the lower surface of the protrusion, the top of the colony was removed to reveal a fixed layer of bacteria below the mobile liquid suspension (Video S2). Unfortunately, the liquid prevented detailed analysis of this lower layer, and flattening the sample left no observable structures, unlike when the pellicle was flattened. The lower biofilm layer therefore lacks the mass and structure of the upper pellicle.

**Fig. 5.**
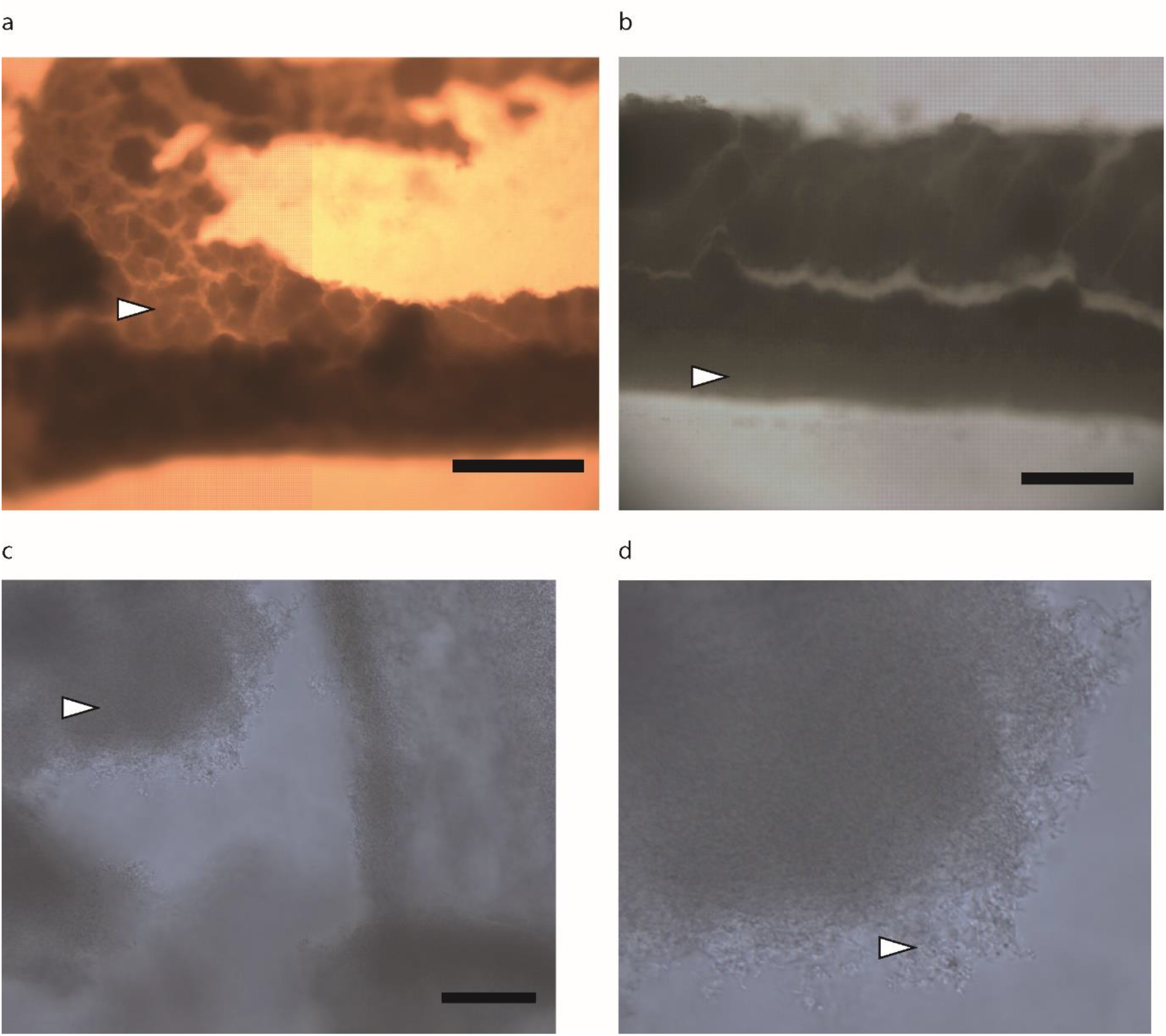
Images of the pellicle enclosing the protrusions of digitate colonies. (a) Fragment of pellicle representative section indicated by the arrow; at the bottom it has folded over). Scale Bar: 500 µm. (b) Another part of the pellicle showing a protrusion edge, the edge has a different consistency to the rest of the pellicle (arrow). Scale Bar: 500 µm. (c) Pellicle fragment showing the edge consisting of disorganised tangled bacteria, the pellicle itself appears relatively thick (representative section indicated by arrow). Scale Bar: 100 µm (d) Enlarged section of (c) showing it is many layers thick and where it is exposed the bacteria formed disorganised but dense arrangements.

To the authors’ knowledge fluid-filled colony protrusions have not been reported previously for any mycobacteria or gram positive bacteria. The closest examples in the literature are some strains of *Roseobacters* (gram negative), which can form liquid channels, but the mechanism for this phenomenon has not been analysed (Bartling *et al*., 2018).

### Fluid expansion in the core of the digitate colony protrusions drives motility

The reduced amounts of medium necessary to permit the use of an inverted microscope tended to result in smaller round colonies. Having established suitable conditions, the circular colonies moved very much as described previously (Video S3). We did not observe growth of any bacteria over the 12 h during which the front of the colony was imaged, and the bacteria did not move relative to each other but were pushed out from within the colony. This behaviour is assumed to arise from unobserved growth at the centre of the colony pushing the bacteria outwards. Martínez *et al*. (1999) also saw no perceptible growth, but by visually tracking the growth of GFP-tagged bacteria they showed the bacteria engaged in 4-6 doublings after 2 days and concluded growth was the basis of the expansion of the circular colonies.

For the digitate colonies, lowering the medium (below 2 ml in the glass-bottomed dish) resulted in more protrusions forming; however, these became smaller with less central liquid, eventually becoming undetectable, although the points of the protrusions remained. The digitate colonies moved in an unusual fashion (Video S4). The protrusions visually expanded by increasing the length of the fluid-filled channel. Also, no growth of the aggregates of bacteria in the fluid core was observed during the period of the experiments (12 h). As the fluid core expanded it progressively pushed aside groups of the bacteria at the tip. When these bacteria were pushed to the sides, they became condensed and aligned to the central core. We provisionally concluded that the fluid expansion itself was the major motive force because bacterial growth has been shown to occur at such a slow rate. In conventional sliding the bacteria can be seen in direct contact with each other and bacteria are being pushed across the whole field of view away from the centre of the colony. From this it is assumed that the force is being transferred from the centre of the colony outwards through the bacteria. This was not observed for the protrusions, where the bacteria were pushed away from the central fluid core rather than the centre of the colony; we therefore assumed the fluid core is the main generator of force in this context.

### New protrusions emerge from sites of physical damage

Damaging the tips of the protrusions resulted in the emergence of fluid, which was at least partially immiscible and did not quickly merge with the agar (Fig. 6a); water spotted on the agar dissipated more rapidly. The damaged tip acted as a new origin for multiple protrusions (Fig. 6b). Disruption of the edge of the protrusion resulted in a bleb, and if heavily damaged a protrusion began to develop from the lesion (Fig. 6b, break shown in Fig. 6c). The pellicle on top of the protrusions could be removed whilst the fluid was retained and still flowed backwards and forwards within the core channel (Fig. 6d, Video S2). After removal the pellicle grew back within 24 h with no obvious scar. Furthermore, removal of the colonies revealed that the bacteria had formed a V-shaped cleft beneath the protrusion that extended to very near the bottom of the plate (Fig. 6e).

**Fig. 6.**
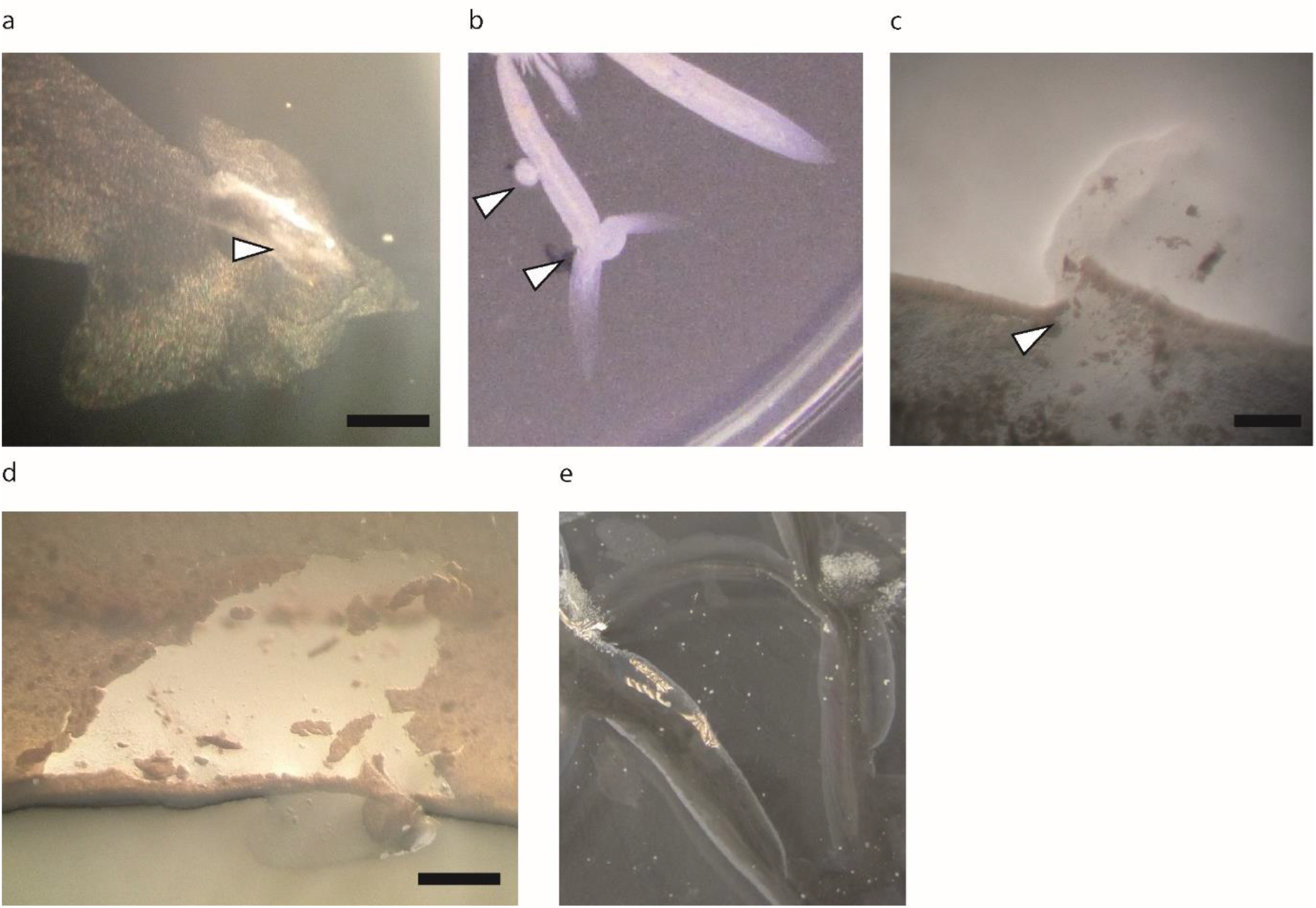
Disruption of a digitate colony. (a) Disrupting the tip of a protrusion results in fluid release (arrow indicates the centre of disruption site). Scale Bar: 1 mm. (b) Letting the colony regrow after disrupting it at the tip results in new protrusions as if starting a new colony (bottom arrow) but only a bleb forming if done in the middle of the protrusion (top arrow). (c) Disruption of the side of a protrusion shows that there is a layer of cells retaining the fluid (arrow site of nicking the edge, note the specialised biofilm edge of the protrusion can be seen clearly here). Scale Bar: 1 mm. (d) The pellicle on top of the protrusion can be removed, leaving both the central core and the edge where it comes in contract with the agar intact. The pellicle can be seen clearly here due to the light angle used in the dissecting microscope. The fluid can still flow through this central core. Scale Bar: 1 mm. (e) When the colony is extracted it leaves V-shaped grooves in the agar that penetrates to nearly the base of the petri dish.

### Digitate colonies form V-shaped clefts in the agar surface

During preparation for SEM the pellicle detached and the core dissipated. This allowed SEM imaging of the internal face of the V-shaped clefts, within which the protrusions were located. The surface of the agar cleft had been modified compared with the surrounding agar (Fig. 7). When agar is desiccated it becomes fibrous (as seen at the top of Fig. 7a) but this was not true of the inside of the cleft where it appeared to be composed of more aggregative strands that penetrated into the surrounding agar (Fig. 7b,c). Furthermore, there were regions of small disorganized groups of bacteria on the surface, and so we assume these bacteria are what remains of the inner biofilm (Fig. 7d).

**Fig. 7.**
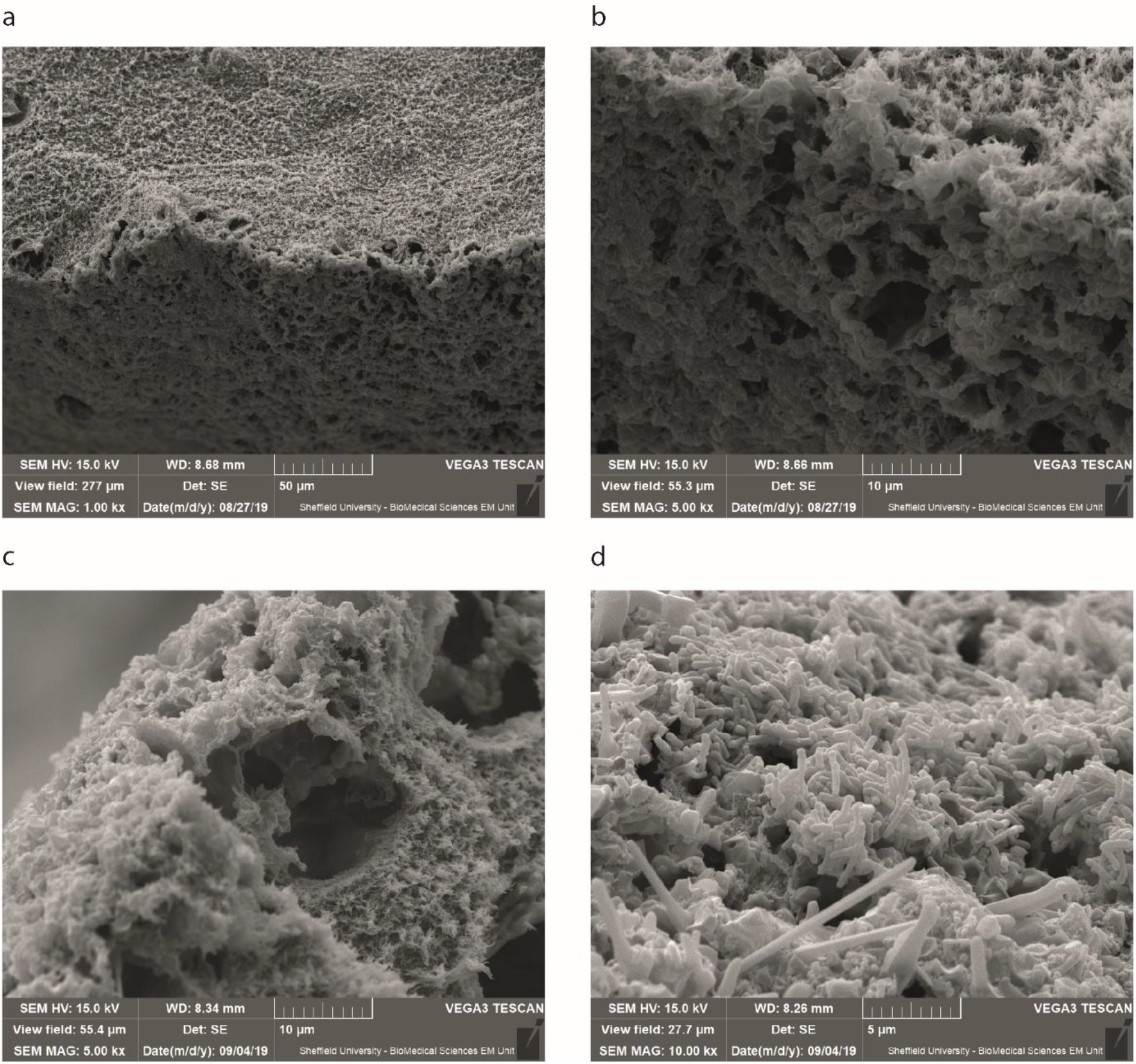
SEM of the inner biofilm. The SEM fixation process removed the pellicle and contents of the protrusions permitting observation of the inner surface (a) The top of the image shows the agar surface, whilst the bottom shows the inside of the protrusion which has a different consistency. (b) The same as (a) but at higher magnification. (c) The edge taken from the opposite angle. The voids are presumed to be where water is being lost. (d) A patch of bacteria on the inside of the protrusion.

### Chemical analysis of the channel fluid

We assessed the feasibility of analysing the core channel fluid using IR and mass spectroscopies. This revealed there were substantial chemical differences in the digitate colonies compared to liquid cultures and very likely different compounds were also present (Fig. S1-4). The IR spectroscopy with its different peaks shows different bond arrangements are present. However, no positive identifications of digitate colony channel fluid components could be made as the peaks overlap in the 200-400 cm^-1^ region where many different bond types are present (Fig. S1-2). Mass spectroscopy PCA score plots showed that the digitate colony samples were fundamentally different from the liquid culture samples (Fig. S3). Putative matches were initially identified against the *E. coli* metabolite database (<u>ecocyc.org</u>) for the most abundant digitate colony peaks (Table S1) (Keseler *et al*., 2017). Remaining unknown peaks were compared to mycobacteria reference mass lists (Zampieri *et al*., 2018). Some of these putatively identified compounds are directly associated with the mycobacterial cell envelope such as p-HBAD (associated with the phenolic glycolipids, a major component of the cell envelope) and Biotinyl-5-AMP (biotinylation being required for the Mycobacterial cell wall). The SDS-PAGE analysis did not reveal major differences in protein content between the *M. smegmatis* liquid culture and the digitate colony extract with a prominent species of ∼150-200 kDa being detected in both samples (Fig. S4). Nevertheless, we have shown that the digitate colony channel fluid can be readily extracted and chemically characterised, which should enable further studies to identify the major components of the channel fluid.

## Discussion

### Mycobacterium smegmatis *forms digitate colonies with centrimetre-long protrusions that facilitate a novel form of sliding motility*

*M. smegmatis* grown on 7H9 medium solidified with Noble agar generated colonies with novel fluid-filled centrimetre-long protrusions. These protrusions consisted of the following components: a central bacterial suspension at the core; retaining biofilm edges in contact with the agar; a surface pellicle; and the tips (Fig. 4-6). The expansion of the central core enabled lengthening of the protrusion, suggesting a novel mechanism of sliding motility as the bacteria at the tip are moved passively by the expansive forces generated by the liquid core, and not directly in response to pushing by growth of bacteria behind as occurs in classical sliding motility (Video S4). This is distinct from the dendrite motility previously described by Martínez *et al*. (1999) where no liquid channel or enclosing biofilms were observed. In conventional sliding motility, bacterial growth generates a circular colony and the bacteria physically push each other outwards through direct contact (Martínez *et al*., 1999; Video S3). The protrusions reported here are therefore radically different as they use a contained liquid component to provide this force. The mechanism by which the mycobacteria draw in and retain fluid in the central channel is unknown, but the fluid must be generating the expansive force as it was observed physically pushing the bacteria forward, moving them to the side and compressing them (Video S4). In conventional sliding colonies, bacterial growth generates the expansive force as the bacteria are pushed radially out from the centre of the colony, maintain their relative position and are usually in direct physical contact with each other. Here instead the bacteria at the front of the colony were pushed to the sides of the protrusion as the fluid pushed forwards. Given these differences we have therefore termed this new form of sliding, *hydraulic sliding*, to reflect its different nature. Under the conditions used, hydraulic sliding could allow colony expansion that is faster than that allowed by the slow growth rate of mycobacterial species. Notably, the closely related *Streptomyces* can engage in a specialist explorer form to move faster than their growth rate would allow (Jones *et al*., 2017). It remains to be determined why the protrusions formed on Noble agar and not agarose, it could be either the overall electrical charge of agarose that could interfere with motility or alternatively it could be due to the specific activity of agaropectin. Additionally, although the behaviour observed could be very different in *M. smegmatis*’ natural environment (soil/water sources) it could represent a way for mycobacteria to enclose an environment to 1) monopolise resources 2) to exclude external predators with biofilms 3) retain metabolically costly secondary metabolites or 4) create a more beneficial environment for growth.

### Mycobacterium smegmatis *digitate colony structure and how it contributes to the fluid channel*

The fluid that enables colony expansion was retained by the structure of the protrusion. The physical retaining components of the protrusion were: 1) the pellicle; 2) the biofilm-like edges; 3) the V-shaped groove in the agar that extended to the base of the petri dish; and 4) the tip of the colony which was pushed outwards (Fig. 4-7, model of the protrusion cross section Fig. 8). It appears that the liquid core is physically retained by the V-shaped groove, by the biofilm like edges and by the tip of the protrusion, because disrupting them causes the fluid core to leak out. The pellicle is believed to prevent the core from drying out as it can be disrupted without causing the core to leak. The V-shaped channel is intriguing as *M. smegmatis* does not have an agarase to digest the agar and so it is likely to be physically pushing aside the agar and retaining the moisture. The SEM images support this hypothesis, because the agar surface of the channel looks different from the surrounding agar. Additionally water put onto an agar surface dissipates, even without the action of surfactant, so fluid must be being trapped within the protrusions to prevent it dissipating (Jain *et al*., 1991). Liquid channels have been seen in several motile *Rhodobacteraceae* strains (aquatic *α-Proteobacteria*) but the mechanism for their formation has not been investigated (Bartling *et al*., 2018). The structure of the colony with a liquid core enclosed by biofilm appears to be unique. It remains an open question how *M. smegmatis* maintains a fluid component when the default for even motile bacteria is to make as physically dense a colony as possible when expanding on agar.

**Fig. 8.**
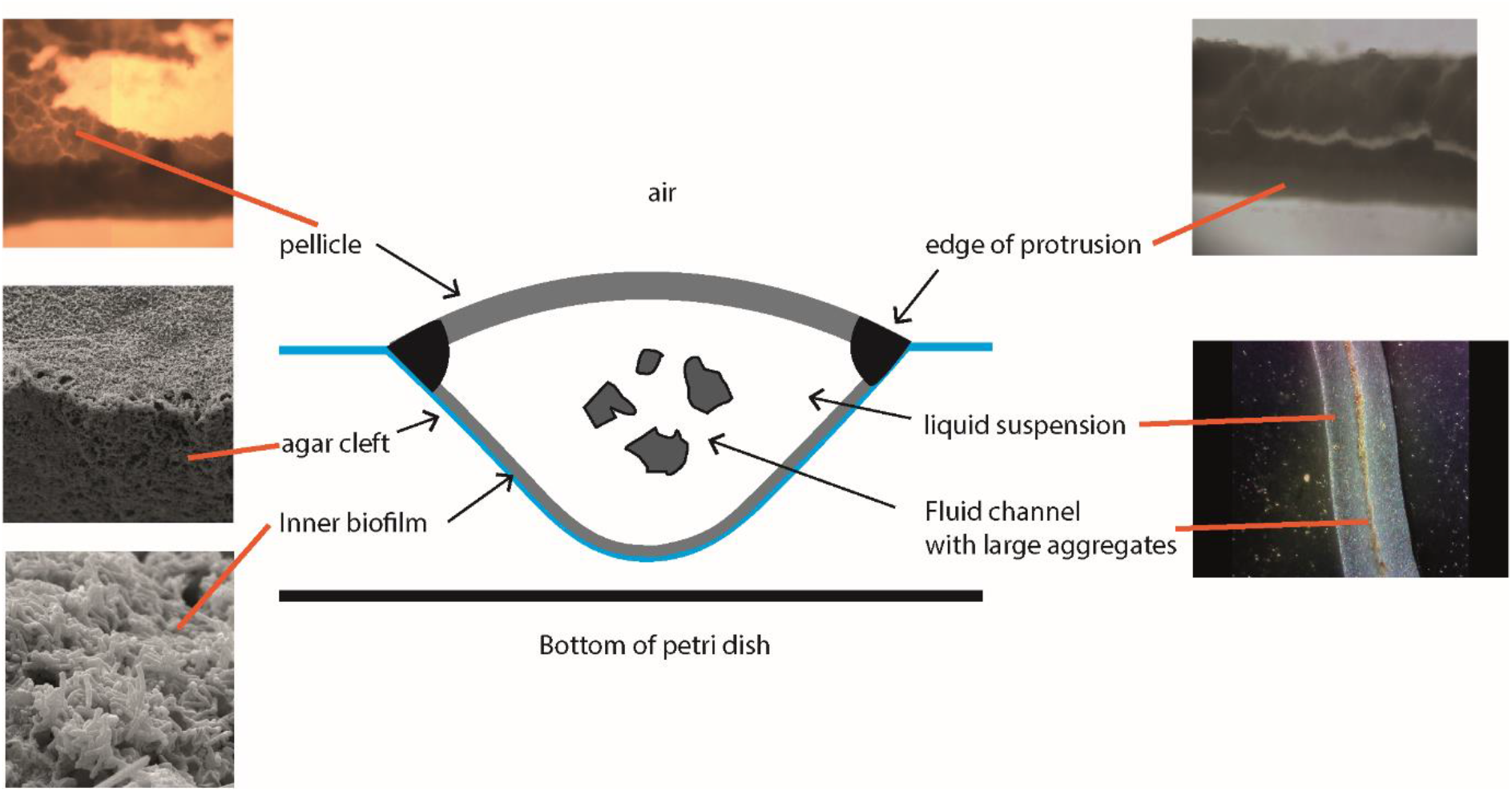
Cross sectional representation of a digitate colony protrusion. A schematic of the *M. smegmatis* sliding protrusion structure. As the fluid suspension/core expands and pushes bacteria to the sides, the protrusion edges build up into specialised structures. Two different types of biofilm form as a pellicle on top of the channel fluid and as a thinner biofilm at the agar/liquid interface. The protrusion structure sits within a V-shaped cleft in the agar that extends almost to the bottom of the petri dish. Within the channel there is a free-flowing core which contains large aggregates of *M. smegmatis*.

### The biofilm components

The fluid within the protrusions is also retained by the surrounding bacteria consisting of a lower layer and the edges in contact with the agar, and the pellicle (Fig. 6). The thin biofilm at the interface of the V-shaped channel represents a very different biofilm type from previously known *M. smegmatis* biofilms (the pellicle and stress-induced biofilm models). Since it does not have any structural strength, it is suggested that it acts to seal the interface with the agar. The pellicle is also different from other biofilm models, in that the bacteria have somehow formed a liquid-air biofilm starting from a solid-air interface and the protrusion pellicle does not appear to wrinkle like ones that form on liquid culture. The pellicle also starts with the branching pseudofilament chains of bacteria rather than the individuals/clumps of bacteria seen with the standard biofilms generated from liquid culture. The edges in contact with the agar have a different hue to the pellicle and are visibly denser than the rest of the pellicle and so are likely to be much more adapted for fluid retention. These novel biofilm structures were not observed for the round *M. smegmatis* colonies, but the round colonies have been shown to require the same components (such as the GPLs) as conventional *M. smegmatis* biofilms (Recht *et al*., 2000). From our results, it would appear mycobacterial biofilms and motility are more closely linked than previously anticipated given the convergence on pellicle formation. There must be an evolutionary reason to generate a structured environment and converge on pellicle formation. Furthermore, in a medical context the BCG vaccine is made from harvested pellicle because it expresses different proteins compared to liquid cultures, implying that pellicle is important in *M. tuberculosis* infection. Further work to determine if *M. tuberculosis* and other mycobacteria also form fluid filled protrusions and the associated pellicles could have interesting implications.

### The chemical and cellular components of the channel

Chemical analysis using IR spectroscopy and mass spectroscopy revealed that the bacteria have a different composition in the digitate colonies compared with those grown in liquid medium using the same nutritional source. Mass spectroscopy found hits putatively associated with components of the mycobacterial cell envelope that occurred more in the digitate colonies than liquid culture. Alterations in the cell envelope could feasibly contribute to the formation of the colony structures seen. Some mycobacteria are particularly notable for cording, a mechanism by which they develop as dense connected cords and which is directly associated with virulence (Kalsum *et al*., 2017). *M. smegmatis* does not have the cording factor required but cording factors have more recently been potentially identified in a wider variety of mycobacteria than previously known (Julián *et al*., 2010). Speculatively, the dense biofilms could be caused by some related cording behaviour or specialised clumping. The GPLs although not bonded to the mycobacteria are considered to be generally retained at the cell surface so are unlikely to be relevant to the fluid but could contribute to the biofilm (Recht *et al*., 2000). It has been noted that mycobacteria produce slime of unknown composition when engaged in sliding, but this tends to be found all over the colony surface and not within the colony (Martínez *et al*., 1999; Arora *et al*., 2008). The inherent hydrophobicity of the mycobacterial cell envelope may also play a part in retaining water in the channels. This suggests multiple distinct ways in which mycobacteria could contribute to the unusual protrusion structure and function. If there are novel products involved (particularly in the fluid) then this could prove very interesting, as products required for movement are often important virulence factors or act as antibiotics (Kearns, 2010; Otto, 2014; Hölscher and Kovács, 2017). An outstanding question is whether *M. smegmatis* secretes substances or polymers during hydraulic sliding; we would assume so given the presence of pellicle and the well-studied role of secreted products in their formation, but this remains to be determined. The free moving aggregates within the channel could resemble those that are seen within liquid culture where this aggregation behaviour is attributed to the relative availability of carbon and nitrogen sources (DePas *et al*., 2019).

The branching pseudofilament groupings of bacteria seen in both colony types are fascinating as they are more like the branching hyphae/vegetative mycelium (which form to penetrate growth substrates) of the closely related *Streptomyces* than the aggregates and single cells usually seen in liquid culture (Scherr and Nguyen, 2009). It is remarkable that they have not previously been studied in detail as they raise several questions; for instance, on a cellular level the formation of branching pseudofilaments in other bacteria is associated with altered cell division and may be occurring here (Flärdh and Buttner, 2009); on a physical level a branching shape is generally poor for moving over surfaces, raising the question as to why this growth form is occurring during mycobacterial sliding motility (Young, 2006).

### Conclusion

This study shows that mycobacteria can engage in a novel form of sliding motility, namely hydraulic sliding. We found that *M. smegmatis* produces fluid-filled centimetre-long protrusions that drive the expansion of the colony. In this case, they can achieve passive motility through the expansion of a liquid channel which pushes the bacteria forwards and outwards forming stable biofilms around the protrusions that emerge. The protrusions may represent a coordinated response to the environment, possibly conceptually similar to how other bacteria are known to form specialised sub-groups and shapes in order to move and form biofilms (Kearns, 2010). There remain many productive avenues of investigation regarding the chemical components of the channel fluid, the mechanical basis of the structure, the new forms of biofilm, and why the bacteria form branching pseudofilaments on surfaces.

## Experimental procedures

### Motility assay protocols

The mycobacterium motility assay with 7H9 medium and *M. smegmatis* strain mc^2^155 described by Martínez *et al*. (1999) had to be modified to permit reproducible behaviour under our laboratory conditions. This was not unexpected due to the widely recognized sensitivity of bacterial movement to specific environmental conditions (Tremblay and Déziel, 2008; Patrick and Kearns, 2009).

A range of concentrations of agarose (0.2, 0.3 and 0.4%) as the solidifying agent was tested to obtain circular sliding colonies. The following protocol yielded reproducible circular colonies. The medium was prepared using agarose (Sigma, A5093; 0.4%) and 7H9 (Oxoid, 1.175 g) in 225 ml of distilled water. This was autoclaved and cooled to ∼55°C. The medium (25 ml aliquots) was then dispensed into 9 cm petri dishes. The plates were left covered in a laminar flow cabinet overnight (not running) to set and then inoculated with colonies taken directly from an *M. smegmatis* stock plate. The plates were then sealed with Parafilm and incubated at 37°C in a humidity-controlled water jacketed incubator (Thermo Scientific Forma 3120). Colonies were visible and started to expand after 2 days, and growth ceased by day 7. It was important to ensure that vibrations were kept to a minimum to maintain the integrity of the plates over the extended time course of the assay.

For the formation of digitate colonies, it was found that Bacto agar (BD), our available equivalent to the Difco agar used by Martínez *et al*. (1999) did not yield satisfactory results (see *Results*). Therefore, a range of concentrations of Noble agar (Oxoid; 0.2, 0.25 and 0.3%) was tested. Based on preliminary experiments the following protocol was adopted. Noble agar (Oxoid; 0.25%) and 7H9 medium (Oxoid, 1.175 g) was added to distilled water (225 ml) and autoclaved. The sterile medium was dispensed into petri dishes, stored, inoculated and incubated as described above.

### Light microscopy

*M. smegmatis* colonies were analysed using a range of microscopic techniques. A benchtop Zeiss microscope was used for observation of bacteria within the colony; a dissecting microscope was used for observing overall colony morphology; and a Nikon Eclipse Ti2 wide field microscope was used for time-lapse recordings of the movement of the edge of the colony. To facilitate time-lapse microscopy, 2 ml of medium in glass windowed mini 35 mm petri dishes (Nunc) permitted digitate and circular colony formation, whilst allowing focussing on the top surface of the agar from below using a Nikon Eclipse Ti2 wide field microscope with a 10x objective. The focal drift compensation mechanism was used to keep the colony in focus during the overnight time-lapse sequence.

In some experiments the colonies were physically manipulated using a 200 µl pipette tip. Where we compressed the samples under the microscope, we first excised a section of the colony by sliding a glass slide underneath and cutting it out of the agar plate with a coverslip. Then whilst viewing it with the microscope (so we could visually track the structures) we put a coverslip over the top and compressed the sample keeping the structures of interest in view. All images are representative of three different independent replicates and microscopy images are shown with the appropriate scale bars.

### Scanning electron microscopy (SEM)

A section of the colony was removed by sliding a glass slide underneath and cutting it out of the agar plate with a coverslip. Sections (2⨯2 cm) were fixed overnight in 2.5% glutaldehyde/0.1M sodium cacodylate buffer. The following day they were washed in buffer, post fixed in 2% aqueous osmium tetroxide and dehydrated in a graded ethanol series and dried in 50% hexamethyldisilazane (HEX) in ethanol. Final drying was in 100% HEX. After removal from the final HEX wash, sections were left to dry overnight, in a fume hood. Samples were mounted onto a pin-stub using a Leit-C sticky tab, gold coated using an Edwards S150B sputter coater and examined in a Tescan Vega3 LMU scanning electron microscope.

### Extraction and processing of digitate colony fluid cores

The fluid cores of the protrusions from multiple colonies (at least 10 combined) grown for 3 days were extracted using a 1000 µl pipette tip, resulting in 20 ml of fluid being available for each analysis. The fluid was freeze dried using a ScanVac Cool Safe 55-4 Pro 3800 for 2 days. For comparison, medium plus agar (20 ml), liquid 7H9 media (20 ml), and liquid *M. smegmatis* culture grown for 3 days (20 ml) were also processed. The samples were analysed by Infrared (IR) and mass spectroscopies, as well as separating any proteins by SDS-polyacrylamide gel electrophoresis (SDS-PAGE).

### IR spectroscopy

The freeze-dried powders were directly analysed using a PerkinElmer Spectrum One Spectrometer with a diamond ATR sampling accessory (eight scans per sample with a resolution of 4 cm^-1^). The powder was removed after measurement using a strong detergent and the plate was cleaned with distilled water before loading the next sample (as the samples tended to strongly adhere to the surfaces and diamond of the spectrometer).

### Mass spectrometry

The solid processed samples were dissolved in a 1:1 mixture of ultrapure water and ultrapure methanol and analysed by directly injecting into a Waters G2 Synapt Mass Spectrometer in both positive and negative ion mode. Peak lists of m/z versus ion counts were generated and manipulated to remove background noise and the data was then binned to 0.2 amu according to the procedure by Overy *et al*. (Overy *et al*., 2005). Principal component analysis plots were generated from these peak lists. The top peaks that were higher in the colony sample and not the control sample were provisionally matched with corresponding masses in the *Escherichia coli* metabolome (ecocyc.org) and mycobcterial component mass matrix (Zampieri *et al*., 2018). Three technical replicates were obtained by analysing the sample three times.

### Protein analysis

Freeze dried samples (see above) were re-suspended in water (100 μl) and mixed with SDS-PAGE loading buffer (1:1), containing 10% (v/v) 2-mercaptoethanol. After boiling for 15 min, the samples were loaded onto a 15% SDS-PAGE gel (with a 4% stacking gel). Polypeptides were separated by electrophoresis at 150 V for 70 min. Polypeptides were stained with Generon Quick Coomassie and imaged on a Gbox Chemi-XX9 imager (Syngene).

## Abbreviations

GPL: glycopeptidolipids
HEX: hexamethyldisilazane
IR: infrared spectroscopy
PSMs: Phenol Soluble Modulins
SDS-PAGE: SDS-polyacrylamide gel electrophoresis
SEM: scanning electron microscopy

## Funding information

This work was supported by White Rose Mechanistic Biology Doctoral Training Programme award (BB/M011151/1) to JG.

## Acknowledgements

We thank the following colleagues at the University of Sheffield for their assistance: Chris Hill for electron microscopy (carried out in the Faculty of Science Electron Microscopy Facility at the University of Sheffield), Heather Walker for the Mass Spectrometry (carried out in the biOMICS Faculty of Science Biological Mass Spectrometry Facility) and Robert Hanson for IR spectrometry; and Simon Foster for the use of laboratory space. We would like to thank Steve Diggle, Phil Elks and Akos Kovacs for providing helpful comments on the manuscript.

## Author contributions

E.J.P., O.C., E.H. and J.G. designed the research; E.J.P. and O.C. performed the experiments; E.J.P., O.C., E.H. and J.G. analysed and interpreted data, and wrote the paper.

## Conflicts of interest

The authors declare that there are no conflicts of interest.

## Notes

### Competing Interest Statement

The authors have declared no competing interest.

