## Supplementary Data for "*Mycobacterium smegmatis* expands across surfaces using hydraulic sliding"

**Fig. S1.** Infrared spectroscopy of liquid culture (red trace) compared with the digitate colony (blue trace). (a) Overall the spectra are similar, but below 2000 $cm^{-1}$ there are dissimilar peaks reflecting the presence of different compounds. Major differences are indicated by arrows. (b) Annotated (wave number) versions of each trace.

**Fig. S2.** (a) Liquid culture bacterial pellet (black trace) compared with the digitate colony bacterial pellet (blue trace) and agar (red trace). There are substantial differences below 2000 $cm^{-1}$ likely indicating different bond arrangements and therefore different compounds. (b and c) Annotated (wave number) versions of each trace.

**Fig. S3**. Mass spectrometry principal component analysis (PCA) score plots. Three technical replicates were used for the analysis. (a) Positive ion mode. (b) Negative ion mode.

**Fig. S4**. SDS-PAGE analyses. Lanes (1) NEB ladder unstained protein standard, Broad Range; (2) blank; (3) 7H9 medium control; (4) *M. smegmatis* liquid culture; (5) *M. smegmatis* digitate colony extract. There is a high molecular weight species in both *M. smegmatis* fractions ~150-200 kDa (arrow).

**Table. S1.** The top metabolite peaks that were higher in the digitate colony samples. Samples marked * have been putatively identified as matching a corresponding mass in the *E. coli* metabolome (ecocyc.org). After this unknown samples marked ** and # were further putatively matched against a Mycobacteria reference mass list [26]. # marked samples can be associated with the Mycobacterial cell envelope. Samples marked – no match was found.

**Supplementary video files**

**Video S1.** Demonstration of fluid mobility of the central core. Rocking the plate backwards and forwards shows how readily the fluid moves within the central core (as can be seen by the aggregates moving within it).

**Video S2.** Colony with the pellicle removed. Rocking of the protrusion with the pellicle removed shows there is a layer at the bottom of the protrusion. The bacterial suspension can be seen moving backwards and forwards as it is rocked over the bottom layer of bacteria.

**Video S3.** Time-lapse microscopy showing expansion of a round colony. Video showing time-lapse microscopy of the expansion of a round colony over the course of 4 h 40 min. The bacteria are pushed forward evenly from behind by the main mass of the colony.

**Video S4.** Time-lapse microscopy showing expansion of a digitate colony. Video showing time-lapse microscopy of the expansion of a digitate colony protrusion over the course of 8 h.

**Supplementary Data**

**Fig. S2. a**

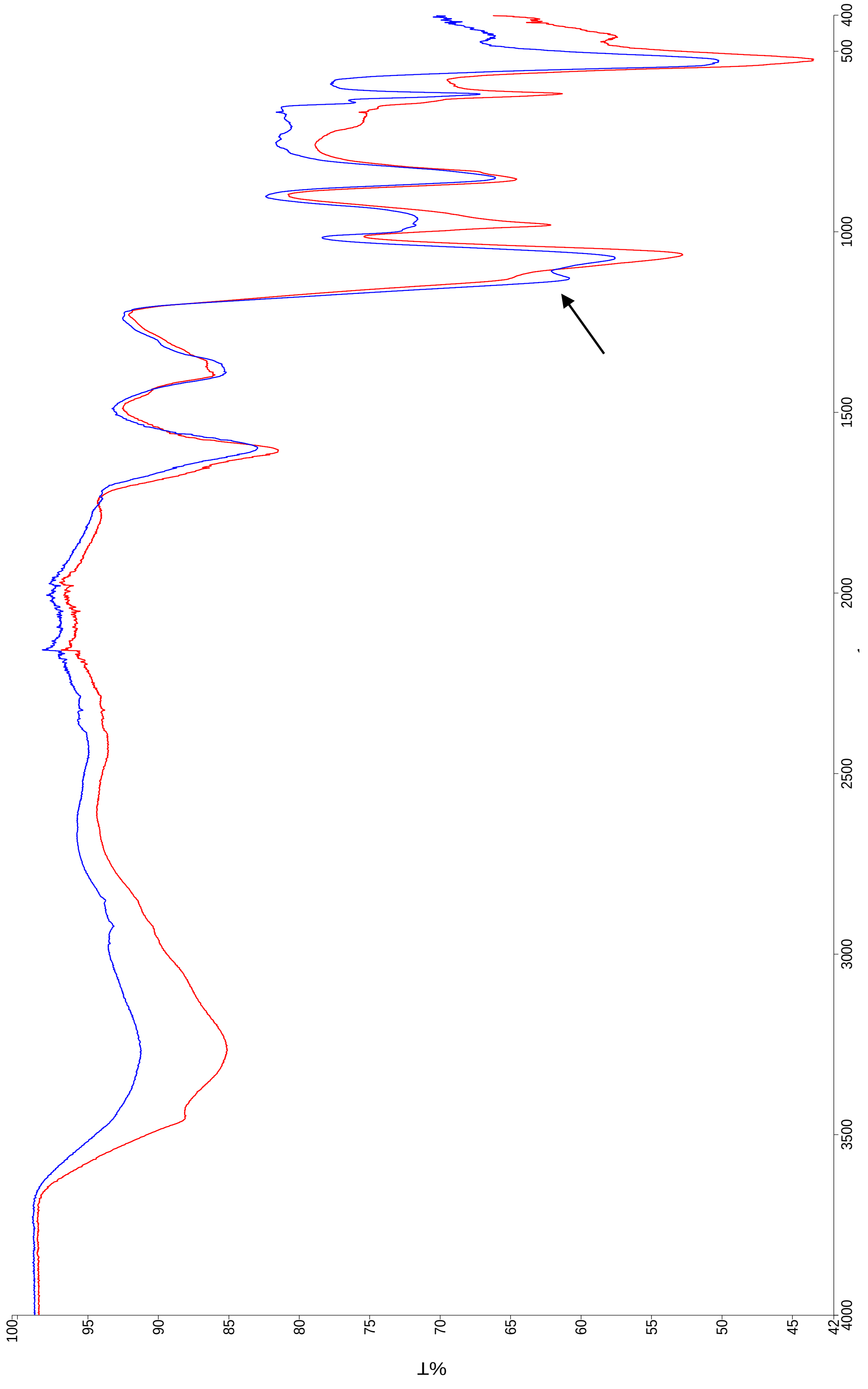

**Fig. S3. b**

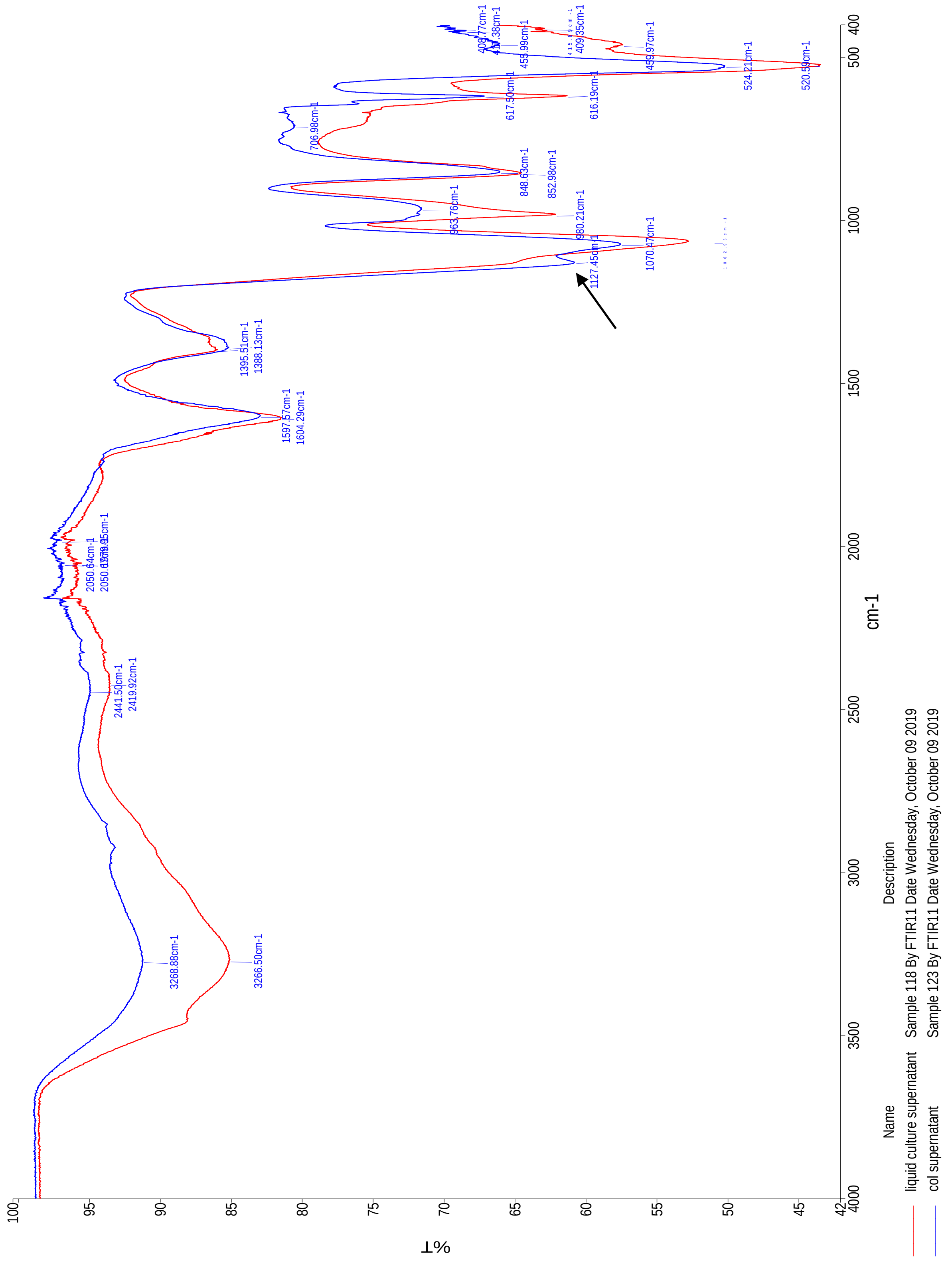
%T
cm-1
Name
Description
liquid culture supernatant
Sample 118 By FTIR11 Date Wednesday, October 09 2019
col supernatant
Sample 123 By FTIR11 Date Wednesday, October 09 2019
3268.88cm-1
3266.50cm-1
2441.50cm-1
2419.92cm-1
1597.57cm-1
1604.29cm-1
1395.51cm-1
1388.13cm-1
1127.45cm-1
1070.47cm-1
980.21cm-1
963.76cm-1
848.63cm-1
852.98cm-1
706.98cm-1
617.50cm-1
616.19cm-1
524.21cm-1
520.59cm-1
459.97cm-1
455.99cm-1
409.35cm-1

**Fig. S2. a**

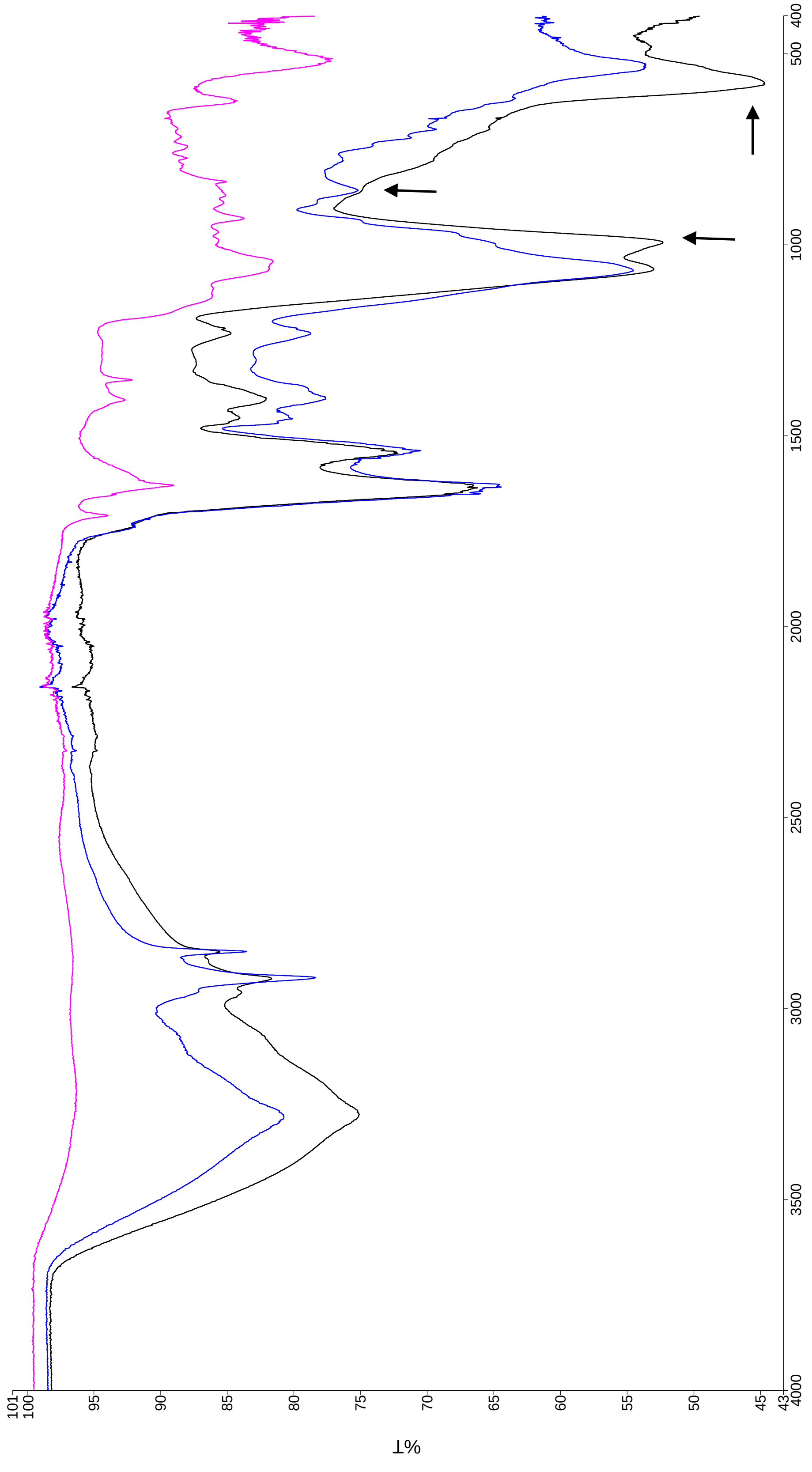

%T
101
100
95
90
85
80
75
70
65
60
55
50
45
43
4000
3500
3000
2500
2000
1500
1000
500
400

**Fig. S2. B**

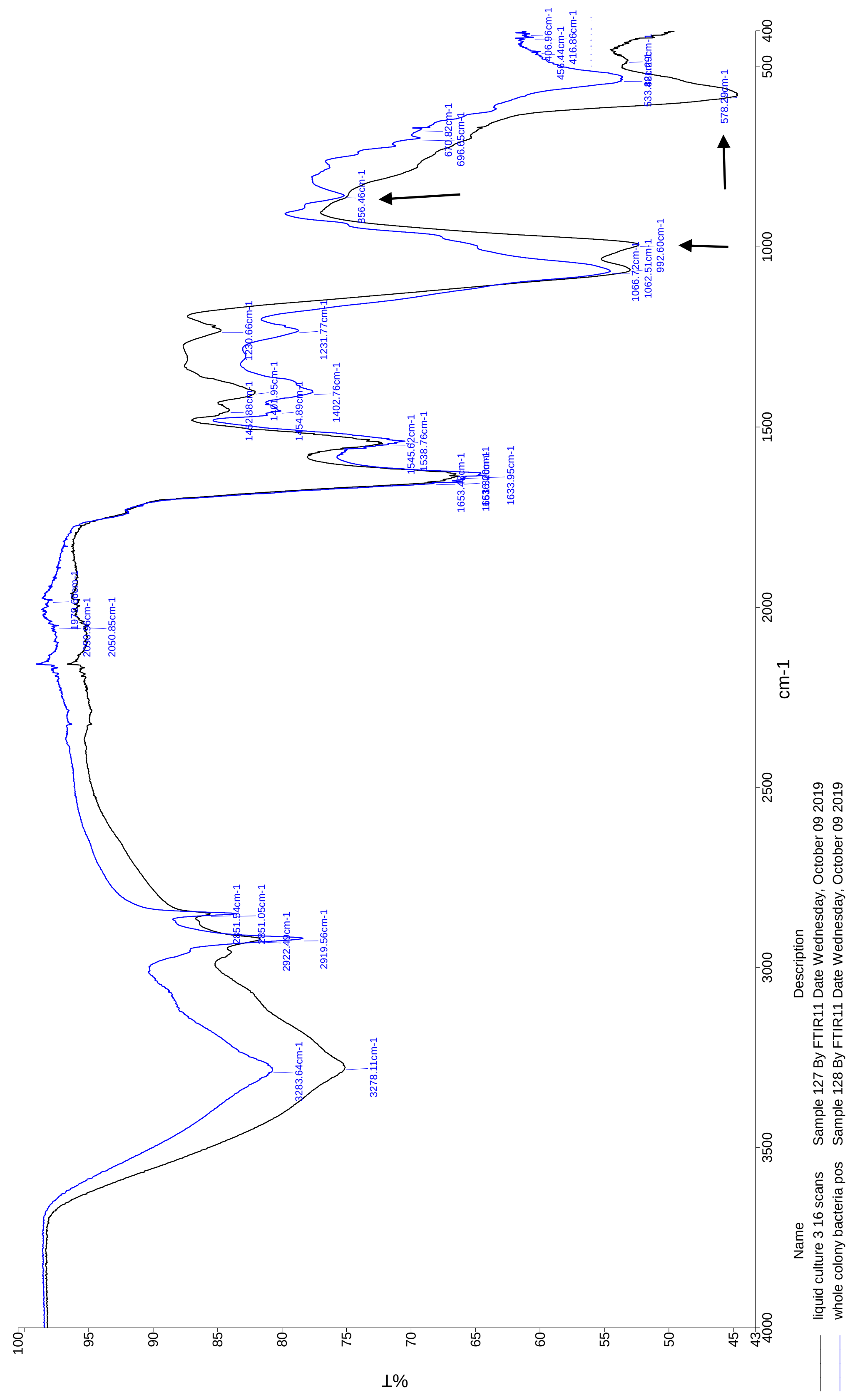
%T
cm-1
Name
Description
liquid culture 3 16 scans
Sample 127 By FTIR11 Date Wednesday, October 09 2019
whole colony bacteria pos
Sample 128 By FTIR11 Date Wednesday, October 09 2019
3278.11cm-1
3283.64cm-1
2919.56cm-1
2922.49cm-1
2851.05cm-1
2851.54cm-1
2050.85cm-1
1633.95cm-1
1545.62cm-1
1538.76cm-1
1402.76cm-1
1231.77cm-1
1230.66cm-1
1062.51cm-1
1066.72cm-1
992.60cm-1
856.46cm-1
696.65cm-1
670.82cm-1
578.29cm-1
416.86cm-1
406.96cm-1

**Fig. S2. c**

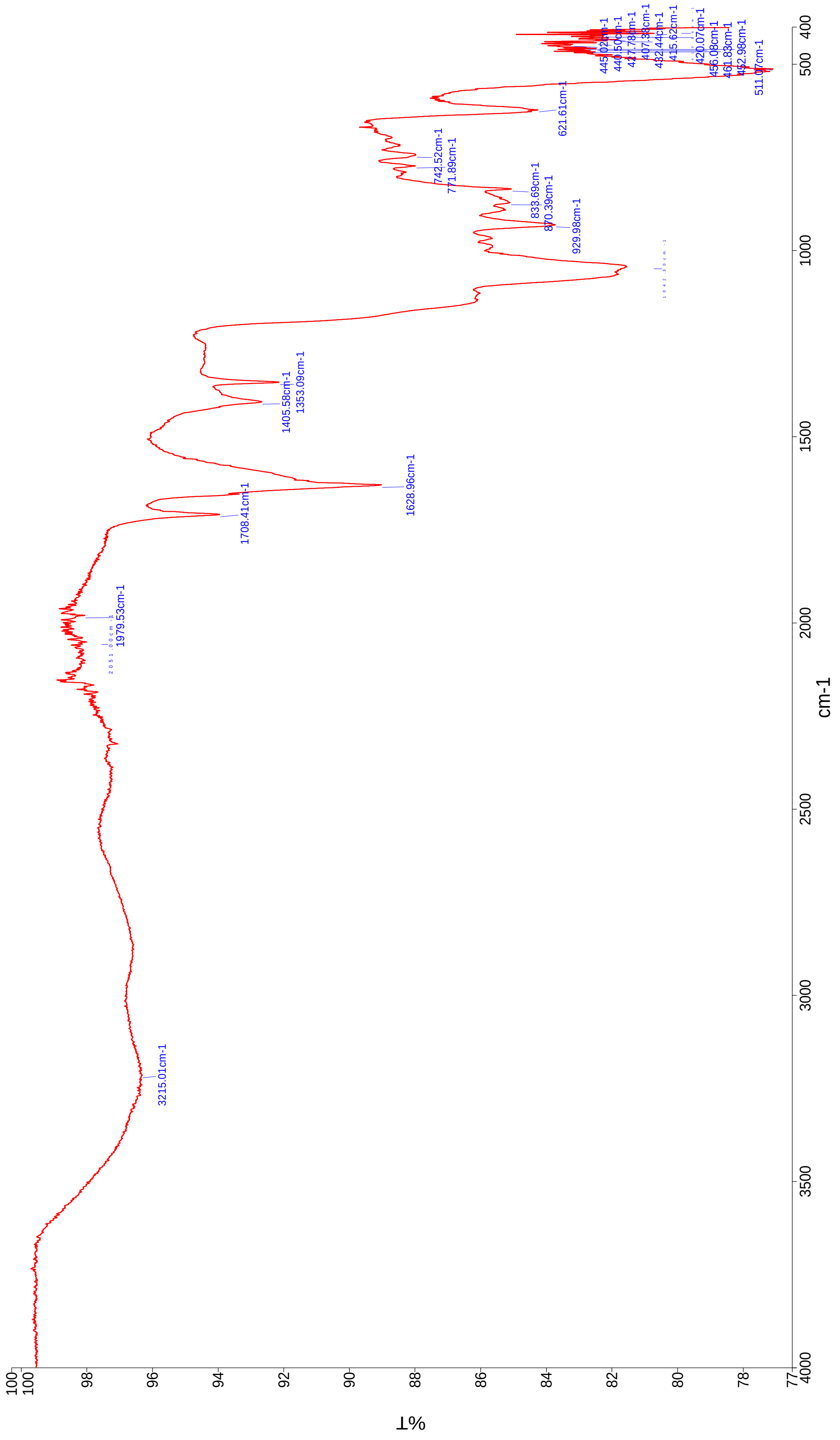

%T
cm-1
3215.01cm-1
1979.53cm-1
1708.41cm-1
1628.96cm-1
1405.58cm-1
1353.09cm-1
929.98cm-1
870.39cm-1
833.69cm-1
771.89cm-1
742.52cm-1
621.61cm-1
511.07cm-1
452.98cm-1
461.83cm-1
456.08cm-1
420.07cm-1
415.62cm-1
432.44cm-1
407.38cm-1
417.78cm-1
440.50cm-1
445.02cm-1

**Fig. S3**.

**a**

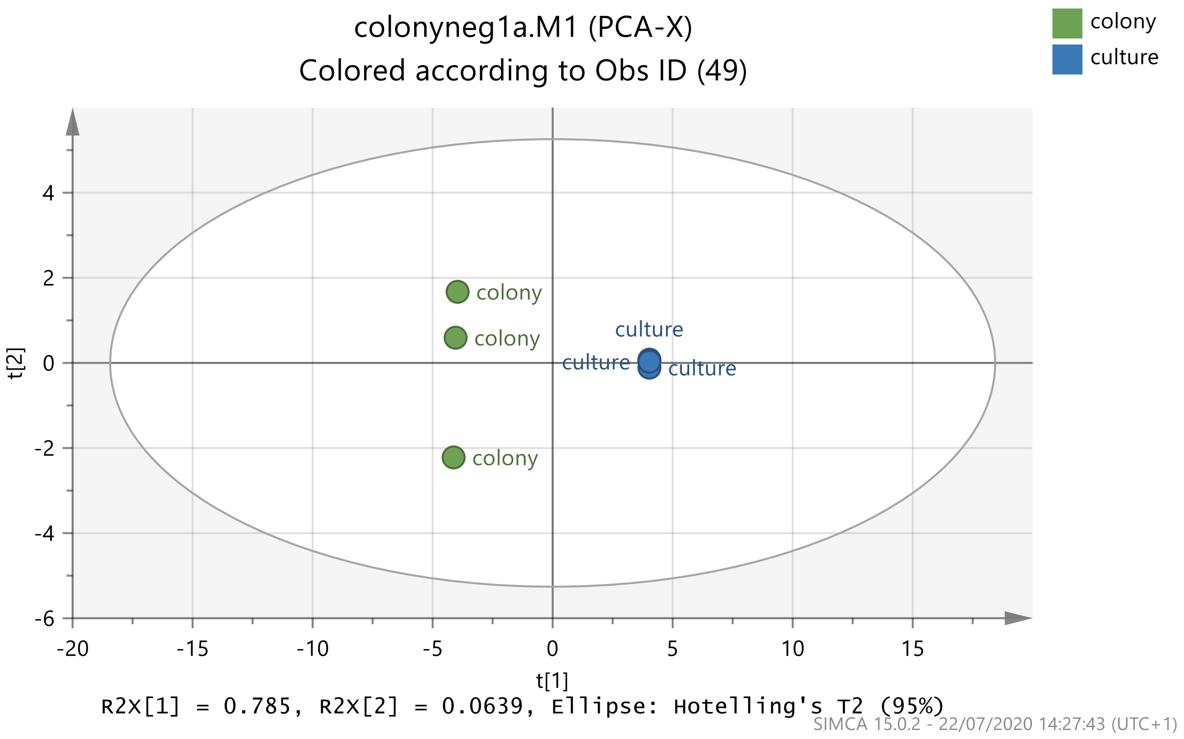
colonyneg1a.M1 (PCA-X)
Colored according to Obs ID (49)
colony
culture
t[2]
t[1]
R2X[1] = 0.785, R2X[2] = 0.0639, Ellipse: Hotelling's T2 (95%)
SIMCA 15.0.2 - 22/07/2020 14:27:43 (UTC+1)

**b**

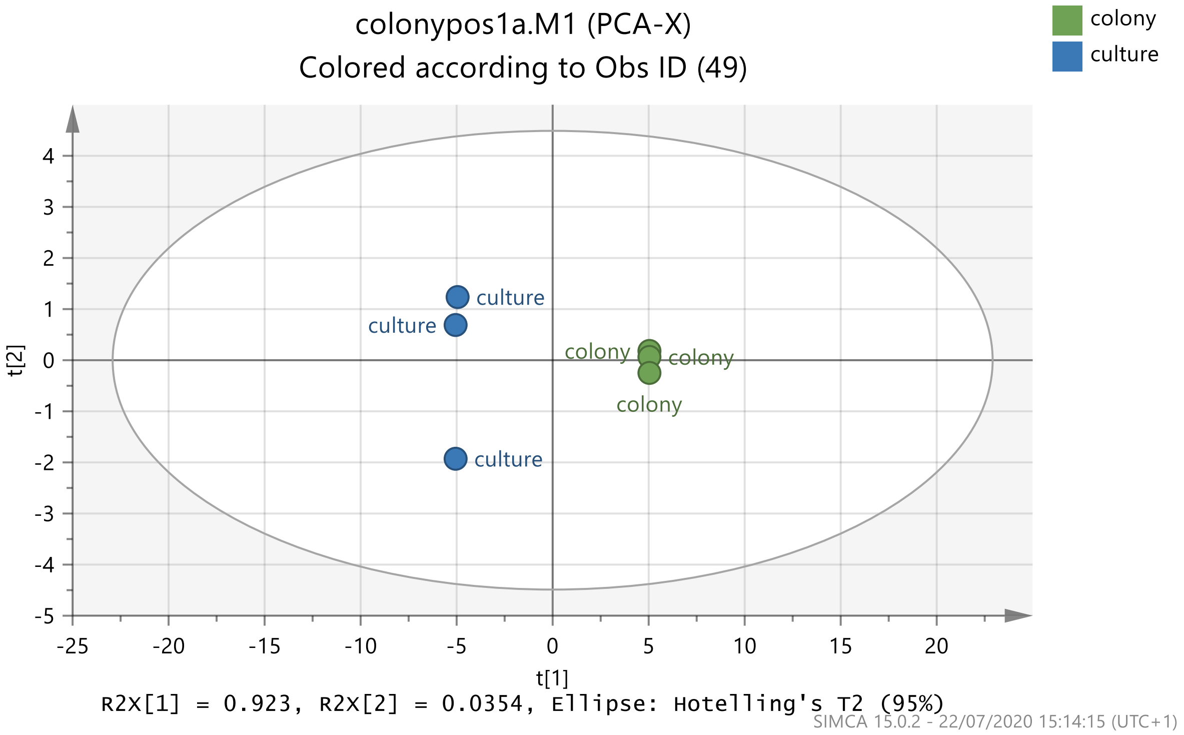
colonypos1a.M1 (PCA-X)
Colored according to Obs ID (49)
colony
culture
t[2]
t[1]
R2X[1] = 0.923, R2X[2] = 0.0354, Ellipse: Hotelling's T2 (95%)
SIMCA 15.0.2 - 22/07/2020 15:14:15 (UTC+1)

**Fig. S4**.

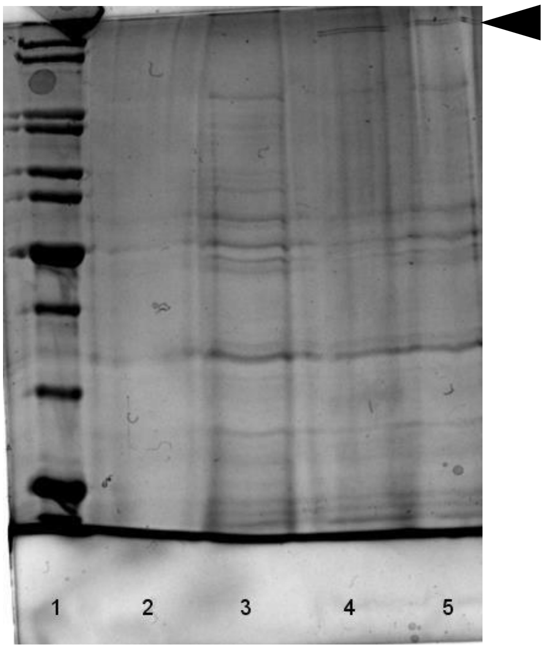

**Table. S1.** The top metabolite peaks that were higher in the digitate colony samples.

| Negative Mode - Bin | Possible ID's |
|---|---|
| 191 | citrate/isocitrate/quinate* |
| 333 | 2-(alpha-D-galactosyl)-sn-glycerol 3-phosphate |
| 282.8 | Guanosine* |
| 221 | cystathionine/flavone* |
| 212 | L-aspartyl-4-phosphate* |
| 712.6 | -2-Octaprenyl-3-methyl-5-hydroxy-6-methoxy-1.4-benzoquinone ** |
| 802.8 | - Phenolpentacosanoyl AMP ** |
| 355 | 5-amino-6-(5'-phosphoribitylamino)uracil* |
| 111 | Uracil* |
| **Positive Mode - Bin** | **Possible ID's** |
| 288.8 | - |
| 430.8 | -N6-Hydroxy-L-lysine coupled to octadecanoyl group# |
| 572.8 | - Biotinyl-5-AMP # |
| 171 | glyceraldehyde-3-phosphate/glycerone phosphate/ornithine/asparagine* |
| 732.6 | - Phosphatidylethanolamine # |
| 448.8 | -Geranylgeranyl diphosphate (GGPP) # |
| 784.8 | - Flavin-adenine dinucleotide-oxidized ** |
| 165 | fucose and hexose deoxy sugar isomers/S-methylmethionine* |
| 590.8 | - Glycosylated p-HBAD I # |
